# Immortalized smooth muscle cells enhance in vitro vasculogenesis

**DOI:** 10.64898/2026.05.08.722734

**Authors:** Mohammad R. Nikmaneshi, Lennard M. Weide, Noel-Adrian Hollosi, Marc Holl, Nemo Nöh, Franciele Filardi Cimino Silva, Dan G. Duda, Lance L. Munn

## Abstract

De novo vessel formation (vasculogenesis) in vitro is a key step in tissue engineering to preserve tissue viability for long-term assays and testing therapeutic agents. However, in vitro vasculogenesis is often unreliable due to differences in vascular-supporting cells, including endothelial cells and stromal cells such as smooth muscle cells (SMCs) and fibroblasts. Here, we developed a robust co-culture system of HUVECs and SMCs to generate stable vascular networks capable of maintaining tissue viability over extended periods. Given that SMC plasticity is a major limitation in supporting endothelial network formation, we systematically evaluated the effects of passage number, confluency, and freezing on primary SMC function. To overcome this limitation, we generated immortalized supportive SMCs, which preserved their vasculogenic gene program and functional capacity even at high passage. In addition, we identified and validated key genes associated with endothelial support, including CD248, C3, and FBLN1, all essential for vasculogenesis. Immortalized SMCs consistently maintained expression of these genes and supported robust vessel formation under variable culture conditions. Collectively, this study demonstrates that immortalized SMCs provide a stable, reproducible platform for endothelial–SMC co-cultures, enabling long-term vascularized tumor models suitable for functional studies and therapeutic screening.

## Introduction

Although current animal models and in vitro cell culture platforms are informative, they have significant shortcomings and inconsistently predict clinical responses [1–4]. For example, standard two-dimensional cultures and spheroids are valuable for identifying targetable pathways in cancer, but they lack essential tumor–stroma–vasculature interactions that mediate tumor growth and treatment response [5–8]. Thus, forming stable vessels in vitro is a critical step for recapitulating cellular interactions and maintaining tissue viability and functionality over extended periods, which is especially important for disease modeling, including cancer[9–13]. Vascular formation generally occurs via distinct mechanisms, including angiogenesis and vasculogenesis[14–16]. Angiogenesis is common in cancers, where new vessels sprout from pre-existing vasculature in response to growth factors produced by tumors[17–19]. In contrast, vasculogenesis, common during development, relies on the de novo assembly of vessels from endothelial cell progenitors, making it particularly suitable for in vitro models[20,21]. Ex vivo vasculogenesis can be used to maintain vascularized tumor explants by allowing endothelial cells to self-assemble into networks from individual cells, thereby enabling greater experimental control over vessel organization in co-culture with tumor tissues or cells[22,23].

Despite these advances, significant challenges remain, primarily due to the high plasticity and senescence of endothelial and vascular-supporting cells, including fibroblasts, smooth muscle cells (SMCs), and pericytes[24–27]. In monocultures embedded in hydrogels such as fibrin, collagen, or Matrigel, endothelial cells can form primitive networks, but these structures are often unstable and require continuous structural support from extracellular matrix–producing cells such as fibroblasts or SMCs[28–31]. Mechanical cues such as shear stress further support vessel maturation, and can be provided through perfusion in microfluidic or 3D culture systems [32–34].

Fibroblasts have been widely used to support endothelial network formation due to their robust ECM production and paracrine signaling that promotes vasculogenesis[35–40]. However, fibroblasts rapidly lose their functional and synthetic capacity after a few passages [36,30,28]. Attempts to immortalize fibroblasts have shown limited success, as immortalized fibroblasts still have a limited functional lifespan, similar to primary cells [41]. Alternative vascular-supporting cells, such as SMCs or pericytes, have been explored. Among ECM-producing cells, SMCs play a key role in vessel formation, angiogenesis, and remodeling due to their dynamic plasticity, which allows switching between contractile and synthetic phenotypes depending on vessel requirements[27,42–47]. This plasticity, however, presents challenges when SMCs are used for vasculogenesis in vitro, as functional support diminishes with passage and culture stress[48].

Relying exclusively on fibroblasts for vessel formation in vascularized tumor models may also introduce confounding effects such as gel contraction. In addition, in cultures containing cancer cells or tumor organoids, cancer-associated fibroblasts (CAFs) may compromise vessel stability[49–52]. Therefore, employing SMCs, which are less abundant in tumors but provide robust vessel support, represents a promising alternative. In our previous studies, we demonstrated that vascular networks co-cultured with SMCs and endothelial cells can preserve tumor viability, physiology, and genomic integrity over extended periods, establishing a reliable in vitro platform for therapeutic testing [43,53].

Here, we report a robust vasculogenesis system using HUVECs and SMCs to generate vascularized tumor models capable of long-term viability and functional testing. We focused on enhancing the robustness of vasculogenesis by maintaining SMC functionality, including generating immortalized SMCs that retain vasculogenic support across higher passages. Our approach addresses the limitations of primary SMC plasticity and provides a reproducible platform for studying tumor–vessel interactions and evaluating anti-cancer therapies in vitro.

## Results and Discussion

In previous work, we demonstrated that an HUVEC:SMC ratio of 1:5 can support vessel formation through vasculogenesis; however, the robustness of this process varies, largely attributable to the condition of the primary SMCs prior to co-culture with HUVECs [53]. To improve reproducibility, we evaluated primary SMCs at their high functional passage (P8) co-cultured with low-passage HUVECs (P2). The results are presented in Figs. 1A and B and correspond to 3 days post co-culture. Our objective was to promote vascular network formation by HUVECs rather than HUVECs clumping. We further optimized the cell ratio based on quantitative metrics and found that a HUVEC:SMC ratio of 1:8 produced the highest vessel density while limiting HUVEC clumping. This ratio was therefore selected for the remainder of the study to minimize fluctuations resulting from either insufficient SMC support or excessive SMC crowding.

**Figure 1.**
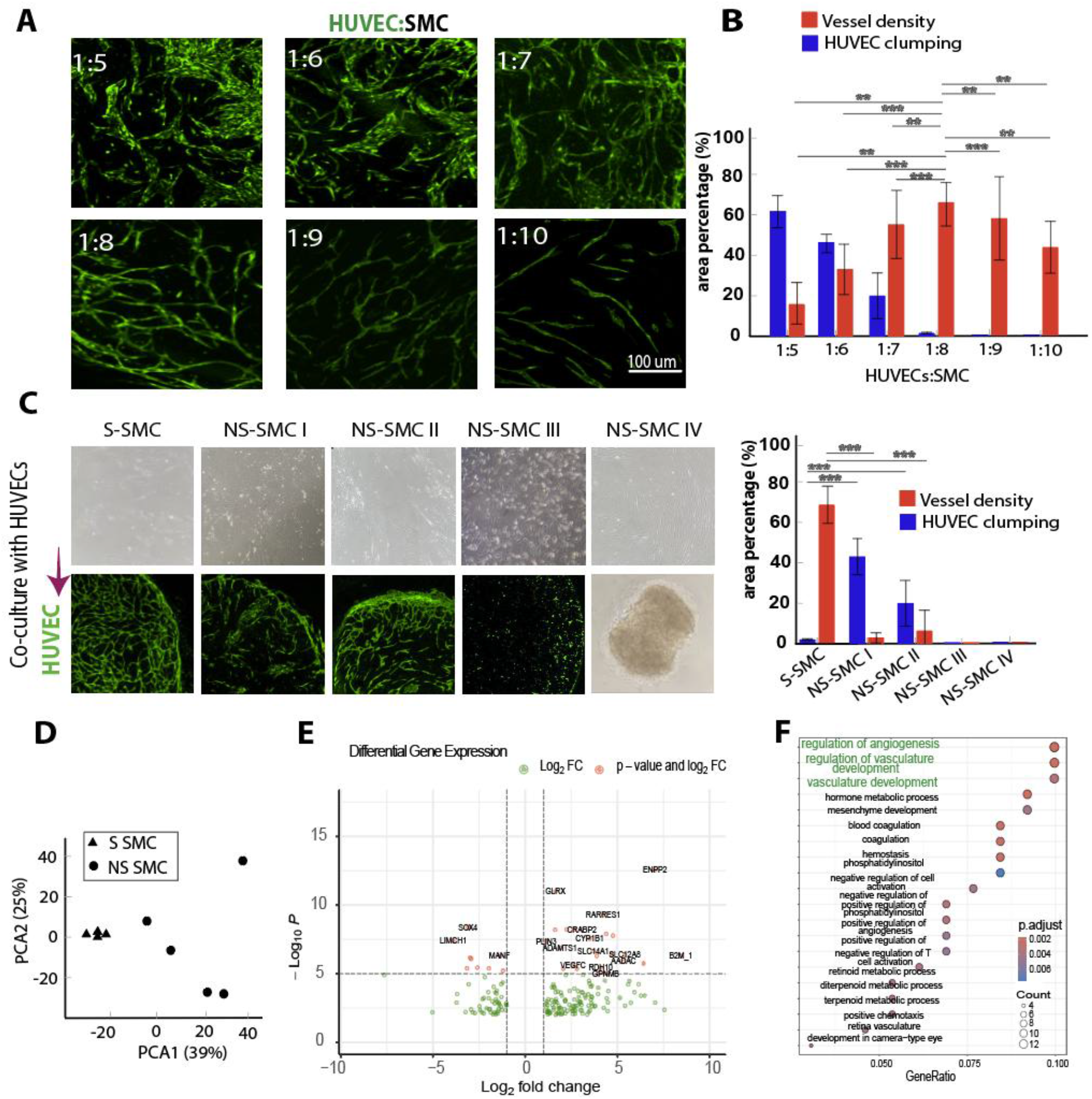
Functional clustering of primary smooth muscle cells (SMCs) based on their ability to support vasculogenesis. Primary SMCs were classified into **supportive SMCs (S-SMCs)** and **non-supportive SMCs (NS-SMCs)** according to their capacity to promote endothelial vessel formation. **(A)** Vasculogenesis assay using co-culture of HUVECs (passage 2) with S-SMCs (passage 8) at different cell ratios, including 1:5–1:10 (HUVEC:SMC). **(B)** Quantification of total vessel density and endothelial cell accumulation as two metrics of vasculogenesis quality. **(C)** Categorization of SMCs into S-SMCs and NS-SMCs based on their ability to support HUVEC vessel formation (SMCs passage 5; HUVECs passage 2). **(D)** Principal component analysis (PCA) of RNA-sequencing data comparing S-SMCs and NS-SMCs. **(E)** Volcano plot showing differentially expressed genes, highlighting the top up- and down-regulated genes based on fold change and adjusted *p*-value. **(F)** Gene Ontology (GO) enrichment analysis of differentially expressed genes, focusing on biological processes.

Another challenge in SMC-driven vasculogenesis is the functional variability among SMC lots. Even lots obtained from the same commercial source and used at comparable passages can yield markedly different vessel-forming outcomes when co-cultured with HUVECs. In an initial screening experiment, we collected multiple SMC lots at a similar passage (P5) and co-cultured them with HUVECs at the same passage (P3). Based on their vasculogenic performance, we categorized them into two groups: “Supportive” SMCs (S-SMCs) and “Non-Supportive” SMCs (NS-SMCs). The monoculture morphology of S-SMCs and NS-SMCs, along with their vessel formation capacity following co-culture, is presented in Fig. 1C. S-SMCs displayed a relatively uniform spindle-shaped morphology with an aspect ratio of approximately 8–10 (SMC mono-culture in the top row of Fig. 1C). In contrast, NS-SMCs exhibited substantial heterogeneity, with aspect ratios ranging from 2 to 20. Functionally, this heterogeneity was associated with variable vasculogenic outcomes, spanning from partial vessel segment formation to complete failure of network formation and, in some cases, the development of a folded or collapsed endothelial network after co-culture. The latter phenotype was likely due to the high contractility of certain NS-SMC populations. NS-SMCs were classified into four groups: NS-SMC I, with an aspect ratio of 8–12 and mean vessel density <5%; NS-SMC II, with an aspect ratio of 10–20 and vessel density <10%; NS-SMC III, with an aspect ratio of 2–5 and no vessel formation; and NS-SMC IV, with an aspect ratio of 10–20, forming no vessels and only a contracted mass.

To investigate the molecular basis of these differences, we lysed all SMC samples shown in Fig. 1C and performed RNA sequencing (RNA-seq) to characterize the transcriptomic profiles of S-SMCs versus NS-SMCs. Principal component analysis (PCA) (Fig. 1D) revealed tight clustering among S-SMC samples, indicating strong transcriptomic similarity, whereas NS-SMCs displayed greater dispersion and were clearly separated from the S-SMC cluster. This separation correlates with the functional divergence observed in Fig. 1C and suggests that S-SMCs share a common transcriptional program, whereas NS-SMCs represent a more heterogeneous population, potentially influenced by distinct underlying factors that impair their supportive capacity.

The volcano plot of differential gene expression (DGE) in Fig. 1E highlights the key genes distinguishing S-SMCs from NS-SMCs. Based on the DGE results, we selected the top 100 differentially expressed genes for gene enrichment analysis. Gene ontology (GO) enrichment analysis for biological processes (Fig. 1F, dot plot) revealed that genes upregulated in S-SMCs were significantly associated with “regulation of angiogenesis,” “regulation of vasculature development,” and “vasculature development.” These findings support the functional classification of S-SMCs and confirm that their transcriptional signature is consistent with a pro-angiogenic and vessel-supportive phenotype.

The ability of primary S-SMCs to support EC vasculogenesis is likely affected by their confluency in monolayer culture before co-culture with ECs and by freeze-thaw cycles. To investigate these effects, we tested batches of S-SMCs at P5 with various confluency levels (70-80 (<80%), 90-100 (>90%), and fully confluent (>100%)). The S-SMCs were co-cultured with HUVECs at P2. The results show that high confluency can reduce the ability of S-SMCs to support HUVECs in forming a network of vessels (Fig. 2A, Fig. 2B). The vascular characteristics, including vessel density, branches per vessel length, average vessel length, and anastomosis (Fig. 2B), demonstrate that S-SMCs at different confluencies exhibit distinct functional states in supporting HUVEC-mediated vessel formation. Thus, for optimal network formation, SMCs should be used when they reach ~90% to 100% confluency.

**Figure 2.**
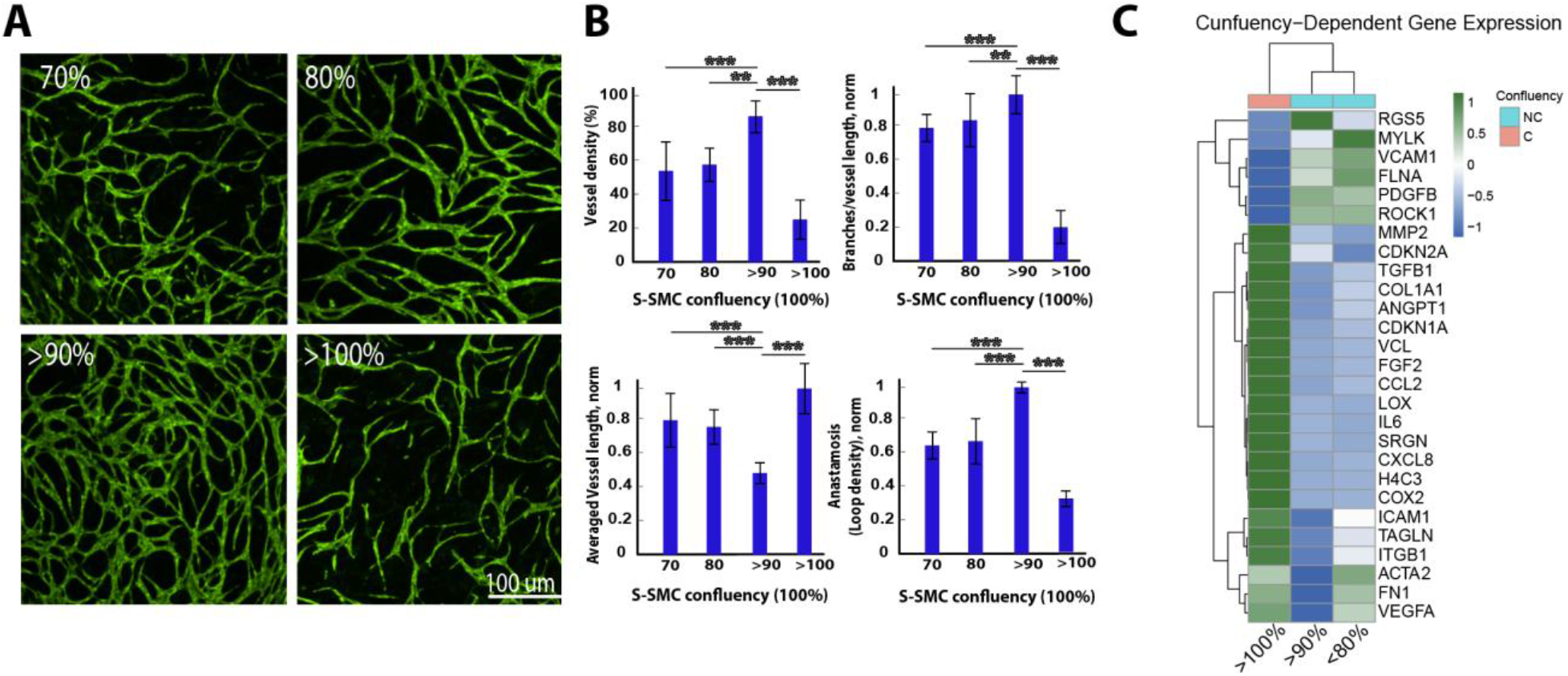
Effect of confluency of SMC on the ability of S-SMCs to support vasculogenesis. **(A)** Representative images of vasculogenesis in 3 day co-cultures with different confluencies of S-SMCs (p5) and HUVECs (p2). **(B)** Comparison of vascular metrics of the cocultures at various S-SMC confluency. **(C)** Analysis of gene expression for SMCs at various confluency levels. Heatmap of row-wise Z-scored VST-normalized expression values across SMC confluency conditions.

The DGE results shown in Fig. 2C for non-confluent (<80% to >90%) vs fully-confluent (>100%) S-SMCs demonstrate that confluency can alter the S-SMC genome. Transcriptomic profiling of S-SMCs under distinct functional states revealed that their ability to support HUVEC vasculogenesis is tightly associated with confluency-dependent shifts in phenotype. The genes of interest comparison between confluency <80%, >90% and >100% (fully-confluent) is shown in Fig. 2C. Slightly subconfluent S-SMCs (>90% confluency) exhibited elevated expression of PDGFB and RGS5, two key regulators of endothelial recruitment and perivascular stabilization[54,55,35], while maintaining relatively low levels of inflammatory mediators such as IL6, CXCL8, and CCL2. This balanced molecular signature is consistent with a supportive perivascular phenotype that promotes organized vessel formation.

In contrast, hyperconfluent S-SMCs (>100% confluency) displayed a pronounced inflammatory and stress-associated transcriptional program. These cells strongly upregulated extracellular matrix and remodeling genes, including COL1A1, FN1, LOX, and MMP2, as well as stress- and inflammation-associated genes H4C3, COX2, and SRGN. Dysregulated pro-angiogenic factors such as VEGFA, ANGPT1, and FGF2 were also elevated, but this activation coincided with reduced PDGFB and RGS5, suggesting potential impairment of endothelial–SMC stabilization. Additional features, including elevated TGFB1, cell-cycle arrest markers CDKN1A and CDKN2A, and mechanotransduction-associated genes such as VCL, indicate a stress-induced, partially senescent phenotype. Collectively, these changes are consistent with a pathological activation state characterized by excessive matrix deposition, inflammatory signaling, and disrupted vascular crosstalk, providing a mechanistic basis for the observed poor vessel formation.

S-SMCs at low confluency (<80%), which exhibit intermediate functional capacity in supporting vascular network formation (Fig. 2A,B) demonstrated increased contractile markers, including ACTA2 and TAGLN, suggesting a shift toward a tension-driven phenotype that may support structural stabilization but lacks the balanced paracrine signaling observed in the optimal condition (>90% confluency).

Overall, these findings indicate that SMC-mediated support of vasculogenesis depends on maintaining a balanced perivascular phenotype characterized by PDGFB- and RGS5-associated endothelial stabilization with limited inflammatory activation. Both suboptimal and overconfluent states disrupt this equilibrium— either through excessive contractility or inflammatory–fibrotic activation—thereby compromising organized vessel formation.

We then used the same S-SMCs at P5 (with confluency >90%), with and without a freeze-thaw (F-T) cycle. The S-SMCs were co-cultured with HUVECs at P2. The results show that F-T can reduce the ability of S-SMCs to support HUVECs in forming a network of vessels (Fig. 3A, Fig. 3B). Key vascular characteristics, including vessel density, branch density per unit vessel length, average vessel length, and anastomosis formation, are modulated by F–T conditions Vascular characteristics, including vessel density, branch frequency per unit vessel length, average vessel length, and anastomosis formation, are modulated by F–T conditions, with F–T cycles reducing the capacity for complete vascular network formation (Fig. 3B). Thus, for optimal network formation, SMCs should be used under conditions of reduced F–T exposure.

**Figure 3.**
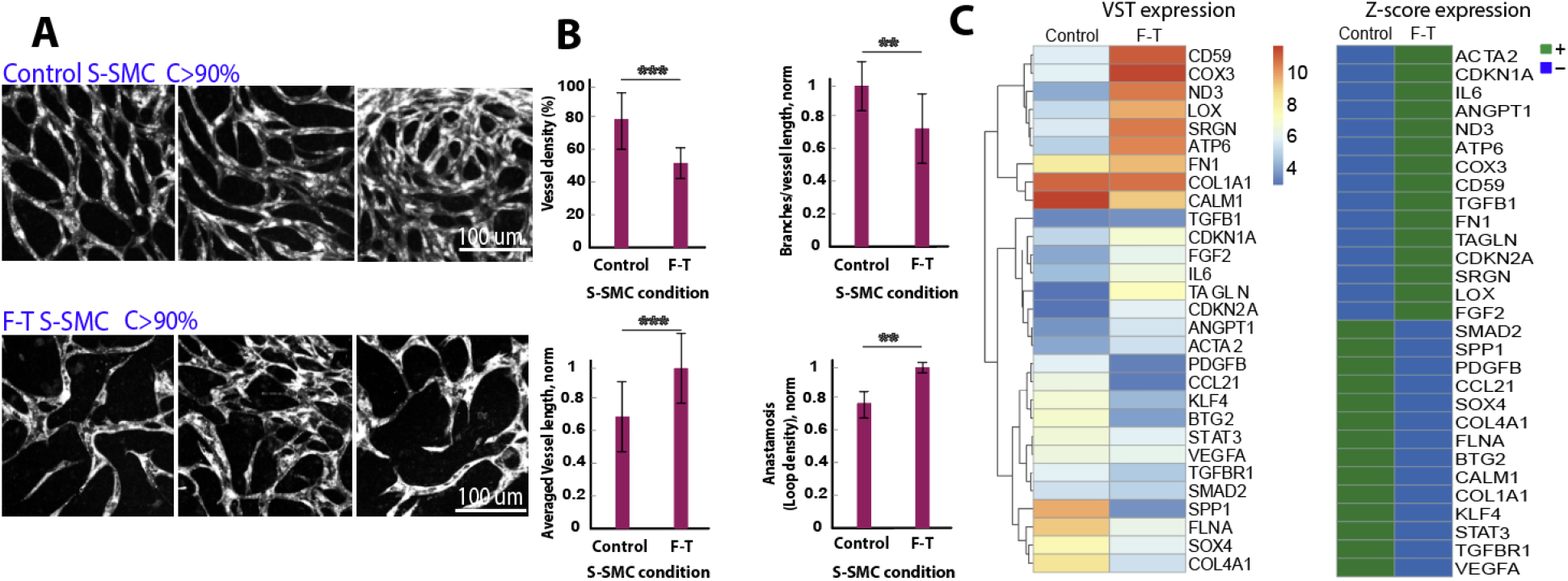
Effect of freezing on the vasculogenic function of S-SMCs. Analyses were performed 3 days after co-culture of S-SMCs (p5) with HUVECs (p2) with and without a freeze-thaw cycle. In all cases, the confluency of SMCs was greater than 90 and less than 100%. (A) Representative images showing vasculogenesis. (B) Quantitative comparison of vascular network metrics across conditions. (C) Gene expression analysis of control and F-T SMCs showing heatmaps of VST-normalized expression values (left), representing relative expression magnitude, and corresponding row-wise Z-scored VST values (right), highlighting relative up- and down-regulation patterns between conditions.

Analysis of Z-scored gene expression in control S-SMCs compared to the same cells exposed to a freeze-thaw (F-T) cycle reveals clear differences in their vasculogenic potential (Fig.3C). Control SMCs exhibit higher relative expression of key angiogenic and ECM-related genes, including VEGFA, PDGFB, FLNA, COL4A1, FN1, SMAD2, KLF4, and STAT3, as well as adhesion and signaling modulators such as SPP1, SOX4, and CALM1. Among these, SPP1, CCL2, PDGFB, and BTG2 are the most upregulated in control compared to F-T SMCs, collectively supporting endothelial migration, proliferation, vessel branching, and stabilization. In contrast, F-T SMCs show elevated expression of stress- and contractility-related genes, including ACTA2, TAGLN, IL6, COX3, and ATP6, with TAGLN and ND3 being most downregulated in control versus F-T cells, indicating reduced metabolic support and diminished capacity to maintain endothelial networks. Some genes with the largest fold changes in control versus F-T SMCs, such as RBM47, RABIF/FIP4, CBR3-AS1, CTSS, and HBE1, may reflect additional aspects of SMC biology but are not directly linked to vasculogenesis. Overall, the Z-score analysis highlights that control SMCs maintain a gene expression program conducive to robust vessel formation, whereas freezing alters their transcriptional profile and impairs functional support for endothelial network development.

Plasticity of SMCs in response to processing effects, such as F-T and confluency, influences their vasculogenic function. A possible strategy to create a more robust system is to use stable, immortalized (Im-) SMCs that keep their vitality and functionality through higher passages.

Comparing Im- and primary S-SMCs co-cultured with the same HUVECs (P2), we found that vasculogenesis is still facilitated by the Im-S-SMCs at high passages, while primary SMCs stop supporting vasculogenesis around passage 8 (Fig.4A). To investigate the mechanism behind the stability of immortalized compared to primary S-SMCs, we performed bulk RNA-sequencing on similar passages P6-8 of primary and immortalized S-SMCs.

**Figure 4.**
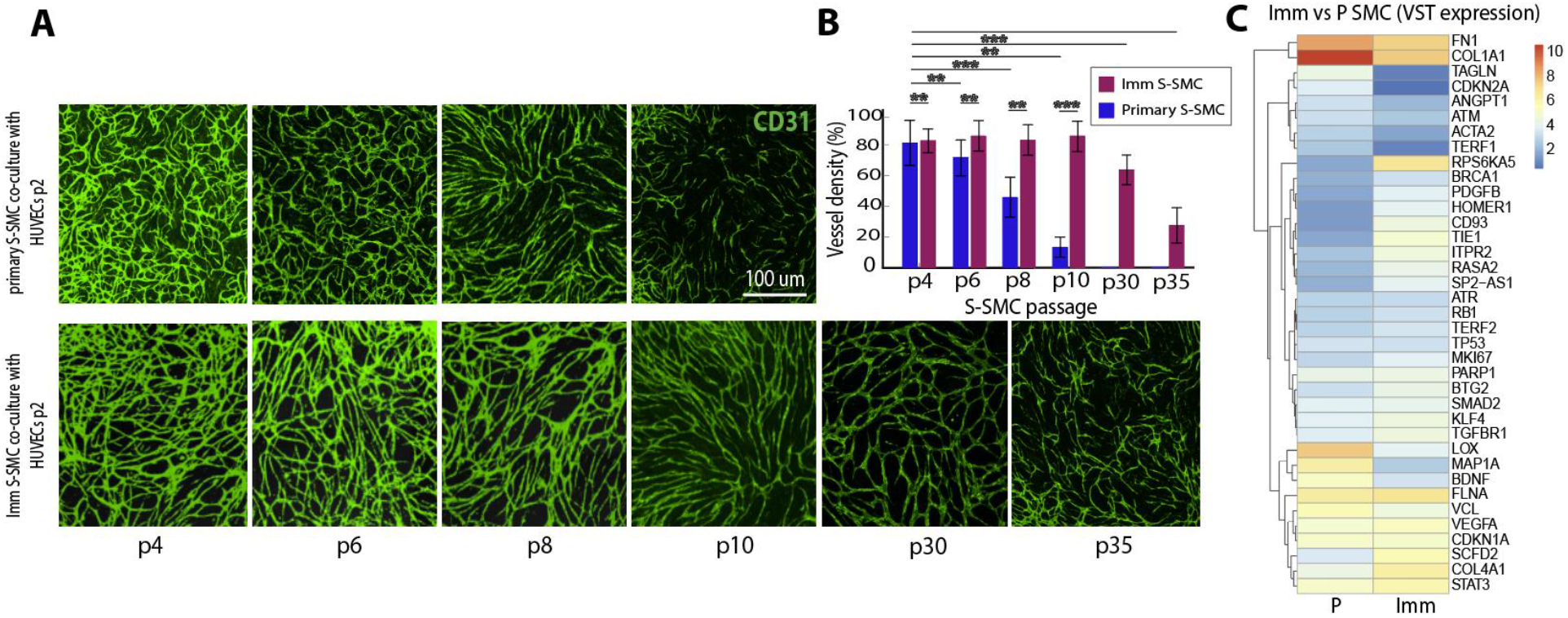
Effect of immortalization of SMCs on vasculogenesis. **(A)** Representative images of vasculogenesis 3 days after co-culture with various confluency levels of S-SMCs (P5) and HUVECs (P2). **(B)** Comparison of vascular metrics. **(C)** VST gene expression in immortalized and primary SMCs.

Analysis of averaged VST-normalized gene expression in early-passage immortalized versus primary S-SMCs reveals that the core vasculogenic program is largely preserved in immortalized cells. Key vasculogenic and signaling genes, including VEGFA, CD93, TIE1, STAT3, KLF4, SMAD2, and TGFBR1, show comparable expression levels between immortalized and primary SMCs, indicating that immortalization does not impair their angiogenic potential.

As expected, genes associated with telomere maintenance and genomic stability, such as TERF1 and RPS6KA5, are elevated in TERT-immortalized S-SMCs, consistent with enhanced cellular stability. TERF1 upregulation likely reflects protective telomere regulation, complementing the proliferative advantage conferred by TERT during immortalization. Structural and cytoskeletal genes, including FLNA and VCL, are slightly higher in immortalized SMCs, which may contribute to stronger adhesion and vessel guidance during vasculogenesis. In contrast, some ECM remodeling genes, notably COL1A1 and LOX, are slightly lower in immortalized SMCs, suggesting minor alterations in extracellular matrix composition; however, this decrease does not appear to compromise overall vasculogenic support.

Collectively, these findings suggest that early-passage immortalized SMCs maintain a functional vasculogenic identity similar to that of primary SMCs, while exhibiting enhanced stability and signaling capacity. This likely underlies their more consistent performance in co-culture vasculogenesis assays and supports their suitability for experimental applications requiring reproducible vessel formation at higher passages.

To further validate the findings of genes of interest associated with SMC support of vasculogenesis identified in the grouping study of S-SMCs and NS-SMCs, we repeated RNA-seq with additional supportive SMC samples. These included both primary S-SMCs and immortalized S-SMCs, generated under controlled conditions of passage (P6-10), non-freezing condition, and optimal confluency (>90%; Fig. 2). These were also compared with primary NS-SMCs used for the grouping study of Fig.1. This expanded analysis improved the ranking of genes identified in the initial study that categorized S-SMCs and NS-SMCs (Fig. 1). Differential gene expression (DGE) analysis for S-SMCs versus NS-SMCs, together with the expression of selected genes involved in the vasculogenic function of SMCs, are shown in Figs. 5A–C.

**Figure 5.**
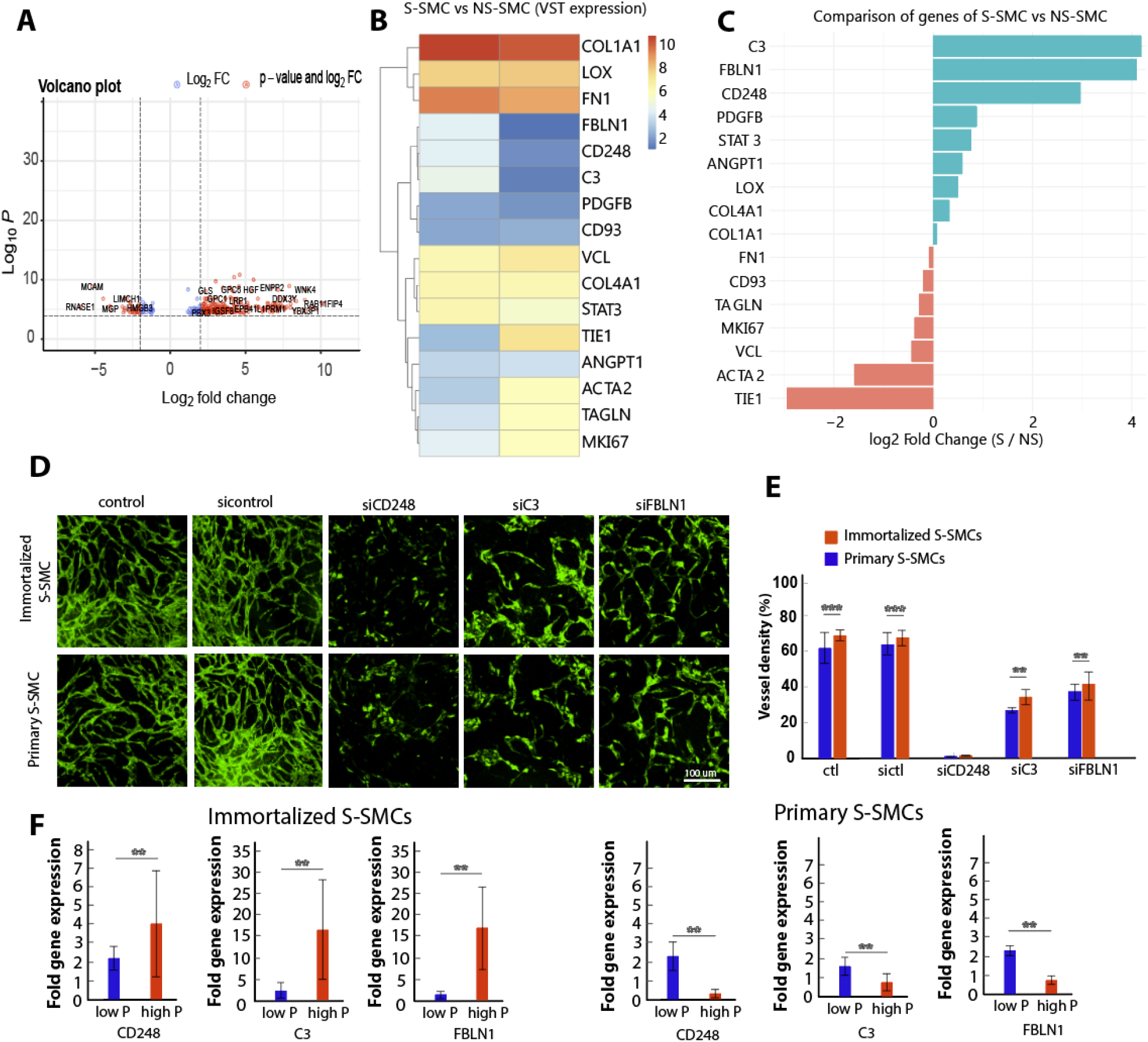
Comparison of supportive SMCs (primary and immortalized) with non-supportive SMCs to identify genes of interest. **(A)** Volcano plot of differentially expressed genes, highlighting the top up- and downregulated genes based on fold change and adjusted p-value. **(B)** VST-normalized expression heatmap of genes related to vasculogenesis in S-SMCs and NS-SMCs. **(C)** Fold-change comparison of selected genes between S-SMCs and NS-SMCs. **(D)** siRNA-mediated functional assay showing the effects of knocking down genes of interest on S-SMC vasculogenic capacity after co-culture with HUVECs (P2). **(E)** Quantification of vessel density from the siRNA assay. **(F)** qPCR analysis of genes of interest in primary and immortalized S-SMCs at low and high passages.

Most of the top genes remained consistent with those previously identified and were associated with vasculogenesis-related processes, although their rankings differed. Among the differentially expressed genes, CD248, C3, and FBLN1 were prioritized for siRNA-mediated functional studies due to their significant regulation in S-SMCs compared with NS-SMCs (Fig. 5C) and their potential roles in perivascular stabilization (CD248), complement-mediated signaling (C3), and extracellular matrix remodeling (FBLN1). These genes were therefore hypothesized to mechanistically contribute to S-SMCs. Based on this consistency, they were selected as markers of the supportive phenotype in both primary and immortalized S-SMCs. We therefore genetically repressed these genes using siRNA and performed functional assays to determine how each affects SMC-mediated vessel formation.

Figure 5D shows vessel formation after 3 days using immortalized and primary S-SMCs (P8, non-frozen, split at >90% confluency) under the conditions: untreated, scrambled siRNA, siCD248, siC3, and siFBLN1. Functional assays demonstrated that CD248 is essential for the vasculogenic support function of S-SMCs, as its genetic repression completely blocked vessel formation. Knockdown of C3 and FBLN1 also impaired vessel formation, though to a lesser extent.

Because functional assays indicated that immortalization preserves SMC function at higher passages (Fig. 4A), we next examined the expression levels of CD248, C3, and FBLN1 across passages. Gene expression was measured in low and high passages of immortalized S-SMCs (low P: 6, 8, 10; high P: 20, 30, 34, 35) and primary SMCs (low P: 5, 6, 8; high P: 10) (Fig. 4B). All three genes remained strongly expressed in immortalized S-SMCs and were even further upregulated at higher passages up to P35, suggesting that they represent core molecular features of the supportive phenotype that are maintained and reinforced during immortalized expansion. In contrast, expression of these genes decreased in higher passages of primary S-SMCs (P10), which may explain why primary SMCs gradually lose their ability to support vessel formation.

Finally, we examined the effects of confluency and freezing on immortalized S-SMCs by comparing gene expression at <80%, >90%, and >100% confluency, as well as in frozen versus non-frozen cultures at comparable passages. Expression of CD248, C3, and FBLN1 remained consistent at high confluency and during freezing, indicating that immortalized S-SMCs maintain the supportive gene program under these conditions. These findings are consistent with functional vasculogenesis assays demonstrating that immortalized SMCs retain vessel-forming capacity even after freezing or growth at high confluency (see supplemental material, Fig. S1).

Our finding that CD248 (endosialin) is essential for S-SMC support of vasculogenesis agrees with studies showing CD248 is upregulated in activated mesenchymal cells and contributes to vascular remodeling and SMC activation in disease models; genetic deletion of CD248 in mice promotes a more contractile SMC phenotype and reduces neointima formation, linking CD248 expression to SMC phenotypic modulation and vascular remodeling in vivo[56–59]. Similarly, FBLN1, an ECM glycoprotein produced by SMCs, has been shown to integrate multiple ECM components and regulate directional SMC migration and tissue remodeling, consistent with our observation that FBLN1 supports vascular network formation likely through ECM organization and cell–matrix interactions[60–64]. Although less studied in SMCs, complement component C3 is part of inflammatory signaling networks that can influence vascular remodeling and SMC behavior via immune–vascular crosstalk[65]; complement activation is associated with vascular inflammation and structural changes that parallel SMC phenotypic shifts reported in other systems[42,65,66]. Collectively, these genes occupy functional niches—perivascular/angiogenic regulation (CD248), inflammatory signaling (C3), and ECM remodeling (FBLN1)—that are intertwined with the plasticity of SMCs previously described in the literature. Our results extend these concepts by functionally linking these markers to SMC support of endothelial network formation, highlighting new mechanistic roles for these genes in regulating supportive versus non-supportive SMC phenotypes within vascular assembly models.

## Conclusion

This study demonstrates that the ability of smooth muscle cells (SMCs) to support endothelial vasculogenesis, beyond dependence on a specific HUVEC:SMC ratio, is governed by a distinct transcriptional and functional phenotype. This phenotype can vary substantially among primary SMC populations and is sensitive to culture conditions such as confluency and freezing as well as passage. We identified key genes associated with the supportive phenotype, including CD248, C3, and FBLN1, and showed that they play distinct roles in regulating vascular network formation.

Importantly, immortalized supportive SMCs preserved the core vasculogenic gene program and maintained robust vessel-supporting function across extended passages. Unlike primary SMCs, whose vasculogenic capacity declines with passage and processing, immortalized SMCs retained stable expression of key supportive genes and remained functionally competent even under conditions that impair primary cells.

Together, these findings indicate that immortalized SMCs provide a stable and reproducible alternative to primary SMCs for endothelial–SMC co-culture assays. Their preserved vasculogenic phenotype and extended lifespan make them a reliable platform for vascular modeling and experimental studies requiring consistent vessel formation.

## Materials and Methods

### Cell Culture

Primary human smooth muscle cells (SMCs) and immortalized SMCs were used to evaluate their ability to support HUVEC vasculogenesis. Primary SMCs were obtained from commercial sources and cultured according to the manufacturer’s instructions. Immortalized SMCs were generated from supportive primary SMC populations and maintained under identical culture conditions to ensure comparability.

SMCs were cultured in Smooth Muscle Growth Medium 2 (SMGM2) supplemented with 10% fetal bovine serum (FBS) and growth supplements, at 37 °C in a humidified incubator with 5% CO_2_. HUVECs were cultured in Vasculife growth medium supplemented with 10% FBS and growth supplements, also at 37 °C in 5% CO_2_. For co-culture experiments, Vasculife medium was used to support both SMCs and HUVECs. HUVECs were maintained at low passage (P2) to minimize phenotypic drift.

When comparing primary and immortalized SMCs, cells were seeded under controlled conditions of passage number, freezing history, and culture confluency (>90%).

For vasculogenesis experiments, SMCs were co-cultured with endothelial cells (ECs). Both ECs and SMCs were first trypsinized, counted, and then mixed at the desired ratio (A:B). In a standard 96-well plate, this corresponded to 10,000 × A HUVECs and 10,000 × B SMCs per well in 200 μL of Vasculife medium.

This setup allowed SMCs to provide structural and paracrine support for endothelial network formation. Co-cultures were maintained for 3–4 days and monitored for vascular network development using phase-contrast microscopy or imaging as appropriate.

To ensure reproducibility, SMCs were cultured under standardized conditions with controlled passage numbers, freeze–thaw history, and confluency (>90% for most assays; <80%, >90%, and >100% specifically for the confluency experiments). Comparative experiments included primary supportive SMCs (S-SMCs), primary non-supportive SMCs (NS-SMCs), and immortalized S-SMCs, which were evaluated for their ability to support endothelial network formation and for differences in gene expression profiles.

### Immortalization of Supportive SMCs (S-SMCs) via hTERT Transduction

#### Determination of Optimal Hygromycin B Concentration (Kill Curve)

To establish stable hTERT-expressing S-SMC lines, the minimal Hygromycin B concentration required to eliminate non-transduced cells was determined:

1. Parental S-SMCs were seeded at 20–25% confluency in a culture plate and incubated at 37°C for 24 hours.
2. Medium was replaced with fresh medium containing a range of Hygromycin B concentrations (50, 100, 250, 500, 750, 1000 μg/mL). Medium was refreshed every 2–3 days.
3. Cell survival was monitored over 7–10 days to identify the lowest concentration that killed the majority of S-SMCs. For these cells, the effective concentration was ~14 ng/mL.

#### hTERT Transduction and Selection

1. S-SMCs were seeded at 300,000 cells per well in a six-well plate. Two wells were prepared for each condition (target and control).
2. The next day, cells were transduced with hTERT virus: 1 mL fresh medium and 1 μL virus enhancement buffer were added to both wells; the target well received 1 mL of virus, and the control well received medium without virus.
3. Media were replaced 24–48 hours post-transduction, and Hygromycin B selection (14 ng/mL) was initiated the following day.
4. Selection continued for 7–10 days, monitoring the control well to ensure complete cell death. Surviving hTERT-expressing S-SMCs were expanded and used for subsequent vasculogenesis assays.

#### RNA Extraction and Gene Expression Analysis

Total RNA was extracted from cultured SMCs using the G-Gene RNA isolation kit according to the manufacturer’s instructions. Cells (~1 × 10^6^) were lysed in 350 μL of RLT buffer, and RNA concentration and purity were assessed using a Nanodrop spectrophotometer. Only samples with high RNA quality (A260/A280 ratio ~1.8–2.0) were used for downstream applications.

For transcriptomic profiling, RNA-seq libraries were generated from high-quality RNA samples (≥30 ng/μL) and sequenced on an Illumina platform. Sequencing reads were aligned to the human reference genome (GRCh38/hg38), and gene-level counts were obtained for differential expression analysis.

Selected genes of interest were validated by quantitative real-time PCR (qPCR). Complementary DNA (cDNA) was synthesized from RNA using reverse transcription, and qPCR was performed with gene-specific primers. Gene expression levels were normalized to the housekeeping gene GAPDH and analyzed using the comparative Ct (ΔΔCt) method.

#### siRNA Transfection – 6-Well Plate Format

##### Aim

To knock down 3 genes using siRNA in human smooth muscle cells (SMCs) with proper controls (untreated and non-targeting siRNA). Cells will be harvested after 72 hours for functional assays and qPCR. Each well should start with ~250,000–300,000 cells.

##### Materials Needed

siRNA (targeting genes; CD248, FLNB1, C3), Non-targeting control siRNA (scrambled), Lipofectamine RNAiMAX (Invitrogen), SmBM (basal media for SMCs), complete SMC media (e.g., SmGM), 6-well tissue culture-treated plates

#### Day 0 – Cell Seeding

1. Trypsinize and count SMCs.
2. Seed 250,000 to 300,000 cells/well in 2 mL of complete SMC media without antibiotics in each well of a 6-well plate.
3. Incubate overnight at 37°C, 5% CO_2_, so that cells reach 60–70% confluence at transfection time.

#### Day 1 – siRNA Transfection

For each well:

##### A. Prepare siRNA–Lipofectamine complexes

- In Tube 1, add 10 μL of 10 μM siRNA to 125 μL of SmBM.
- In Tube 2, dilute 5 μL of Lipofectamine RNAiMAX in 125 μL SmBM.
- Combine both tubes (total ~250 μL), gently mix by pipetting or flicking, and incubate at room temperature for 15–20 minutes to allow complex formation.

##### B. Add complexes to cells

- Remove existing media from cells and replace with 1.75 mL of fresh antibiotic-free complete media (SmGM without antibiotics).
- Add the 250 μL siRNA–Lipofectamine mix dropwise to the cells.
- Gently rock or swirl the plate to distribute evenly.

##### Post-Transfection Incubation

- Incubate cells at 37°C, 5% CO_2_.
- Do not disturb for at least 6 hours.
- After 6–8 hrs (only for primary SMCs not immortalized SMCs): Replace with 2 mL fresh SmGM (to reduce toxicity).
- Continue incubation up to 72 hrs.

#### Day 3–4 – Assays

- Proceed with:
- –Functional assays (Co-culture of 80k SMCs with 10k HUVECs in each well of a 96-well plate.)
- –RNA extraction for qPCR
- –Microscopy

#### Bioinformatics and Differential Gene Expression Analysis

RNA-seq data were processed and analyzed using standard bioinformatics pipelines. Raw sequencing reads were quality-controlled, aligned to the human genome, and quantified to obtain gene expression counts.

Differential gene expression analysis between S-SMCs and NS-SMCs was performed using statistical models implemented in the DESeq2 framework. Genes with significant fold changes and adjusted p-values below predefined thresholds were considered differentially expressed.

To identify biological pathways associated with the supportive SMC phenotype, Gene Ontology (GO) enrichment analysis was performed for the following categories: Biological Process (BP), Cellular Component (CC), and Molecular Function (MF).

Enrichment analysis was conducted using publicly available bioinformatics tools and R packages. The top enriched GO terms were visualized using bar plots and clustering approaches.

#### Functional and Imaging Assays

Endothelial network formation was evaluated using co-culture vasculogenesis assays. Endothelial cells were seeded on SMC layers and monitored for network formation over time. Images were captured using phase-contrast or fluorescence microscopy.

Quantitative analysis of vascular networks included measurements such as total network length, branch points, and vessel density using image analysis MATLAB code.

#### Statistical Analysis

All experiments were performed with biological replicates unless otherwise stated. Data are presented as mean ± standard deviation (SD). Statistical comparisons between groups were performed using appropriate tests, including Student’s t-test when multiple groups were compared. Correlation analyses were conducted where applicable. Statistical significance was defined as *p* < 0.05 unless otherwise specified.

## Acknowledgement

Lance Munn’s research is supported by NIH R01CA247441, U01CA261842, R01CA284603 and a grant from the Additional Ventures Foundation (1452100). We thank Mark Duquette, Hajime Taniguchi, Sonu Subhudi, Rieke Schleinhege, and Pinji Lee for their valuable consultations.

## References

1. Matos LO, Sordi MB, Birjandi AA, Sharpe PT, Cruz ACC (2025) Breaking Barriers: Evaluating Challenges in Advancing Periodontal Ligament Cell-Derived Organoids. Dent J (Basel) 13 (9). doi:10.3390/dj13090422

2. Huang T, Huang W, Bian Q (2025) Organoids as predictive platforms: advancing disease modeling, therapeutic innovation, and drug delivery systems. J Control Release 387:114222. doi:10.1016/j.jconrel.2025.114222

3. Harter MF, Recaldin T, Gjorevski N (2025) Organoids as models of immune-organ interaction. Cell Rep 44 (9):116214. doi:10.1016/j.celrep.2025.116214

4. Ghaemmaghami AM, Hancock MJ, Harrington H, Kaji H, Khademhosseini A (2012) Biomimetic tissues on a chip for drug discovery. Drug discovery today 17 (3-4):173–181. doi:10.1016/j.drudis.2011.10.029

5. McAllister SS, Weinberg RA (2010) Tumor-host interactions: a far-reaching relationship. J Clin Oncol 28 (26):4022–4028. doi:10.1200/JCO.2010.28.4257

6. Jiang P, Li X, Thompson CB, Huang Z, Araiza F, Osgood R, Wei G, Feldmann M, Frost GI, Shepard HM (2012) Effective targeting of the tumor microenvironment for cancer therapy. Anticancer research 32 (4):1203–1212

7. Duda DG, Duyverman AM, Kohno M, Snuderl M, Steller EJ, Fukumura D, Jain RK (2010) Malignant cells facilitate lung metastasis by bringing their own soil. Proc Natl Acad Sci U S A 107 (50):21677–21682. doi:10.1073/pnas.1016234107

8. Nia HT, Datta M, Seano G, Zhang S, Ho WW, Roberge S, Huang P, Munn LL, Jain RK (2020) In vivo compression and imaging in mouse brain to measure the effects of solid stress. Nat Protoc 15 (8):2321–2340. doi:10.1038/s41596-020-0328-2

9. Carmeliet P, Jain RK (2011) Molecular mechanisms and clinical applications of angiogenesis. Nature 473 (7347):298–307

10. Ren B, Jiang Z, Murfee WL, Katz AJ, Siemann D, Huang Y (2023) Realizations of vascularized tissues: From in vitro platforms to in vivo grafts. Biophysics Reviews 4 (1)

11. Meng X, Xing Y, Li J, Deng C, Li Y, Ren X, Zhang D (2021) Rebuilding the Vascular Network: In vivo and in vitro Approaches. Frontiers in Cell and Developmental Biology 9:639299

12. Liu Q, Ying G, Hu C, Du L, Zhang H, Wang Z, Yue H, Yetisen AK, Wang G, Shen Y (2025) Engineering in vitro vascular microsystems. Microsystems & nanoengineering 11 (1):100

13. Chen SW, Blazeski A, Zhang S, Shelton SE, Offeddu GS, Kamm RD (2023) Development of a perfusable, hierarchical microvasculature-on-a-chip model. Lab on a Chip 23 (20):4552–4564

14. Patel-Hett S, D’Amore PA (2011) Signal transduction in vasculogenesis and developmental angiogenesis. The International journal of developmental biology 55:353

15. Vailhé B, Vittet D, Feige J-J (2001) In vitro models of vasculogenesis and angiogenesis. Laboratory investigation 81 (4):439–452

16. Poole TJ, Coffin JD (1989) Vasculogenesis and angiogenesis: two distinct morphogenetic mechanisms establish embryonic vascular pattern. Journal of experimental zoology 251 (2):224–231

17. Hillen F, Griffioen AW (2007) Tumour vascularization: sprouting angiogenesis and beyond. Cancer and Metastasis Reviews 26 (3):489–502

18. Liu Z-L, Chen H-H, Zheng L-L, Sun L-P, Shi L (2023) Angiogenic signaling pathways and anti-angiogenic therapy for cancer. Signal transduction and targeted therapy 8 (1):198

19. Zhang R, Yao Y, Gao H, Hu X (2024) Mechanisms of angiogenesis in tumour. Frontiers in oncology 14:1359069

20. Yin H, Wang Y, Liu N, Zhong S, Li L, Zhang Q, Liu Z, Yue T (2024) Advances in the model structure of in vitro vascularized organ-on-a-chip. Cyborg and bionic systems 5:0107

21. Czirok A, Little CD (2012) Pattern formation during vasculogenesis. Birth Defects Research Part C: Embryo Today: Reviews 96 (2):153–162

22. Cheng G, Liao S, Kit Wong H, Lacorre DA, di Tomaso E, Au P, Fukumura D, Jain RK, Munn LL (2011) Engineered blood vessel networks connect to host vasculature via wrapping-and-tapping anastomosis. Blood 118 (17):4740–4749. doi:10.1182/blood-2011-02-338426

23. Gruionu G, Bazou D, Maimon N, Onita-Lenco M, Gruionu LG, Huang P, Munn LL (2016) Implantable tissue isolation chambers for analyzing tumor dynamics in vivo. Lab Chip 16 (10):1840–1851. doi:10.1039/c6lc00237d

24. Wang G, Jacquet L, Karamariti E, Xu Q (2015) Origin and differentiation of vascular smooth muscle cells. The Journal of physiology 593 (14):3013–3030

25. Song H-HG, Rumma RT, Ozaki CK, Edelman ER, Chen CS (2018) Vascular tissue engineering: progress, challenges, and clinical promise. Cell stem cell 22 (3):340–354

26. Kennedy CC, Brown EE, Abutaleb NO, Truskey GA (2021) Development and application of endothelial cells derived from pluripotent stem cells in microphysiological systems models. Frontiers in cardiovascular medicine 8:625016

27. Beamish JA, He P, Kottke-Marchant K, Marchant RE (2010) Molecular regulation of contractile smooth muscle cell phenotype: implications for vascular tissue engineering. Tissue Engineering Part B: Reviews 16 (5):467–491

28. Kim S, Lee H, Chung M, Jeon NL (2013) Engineering of functional, perfusable 3D microvascular networks on a chip. Lab on a Chip 13 (8):1489–1500

29. Au P, Tam J, Fukumura D, Jain RK (2008) Bone marrow–derived mesenchymal stem cells facilitate engineering of long-lasting functional vasculature. Blood, The Journal of the American Society of Hematology 111 (9):4551–4558

30. Koh W, Stratman AN, Sacharidou A, Davis GE (2008) In vitro three dimensional collagen matrix models of endothelial lumen formation during vasculogenesis and angiogenesis. Methods in enzymology 443:83–101

31. Davis GE, Senger DR (2005) Endothelial extracellular matrix: biosynthesis, remodeling, and functions during vascular morphogenesis and neovessel stabilization. Circulation research 97 (11):1093–1107

32. Pries A, Secomb T, Gaehtgens P (1998) Structural adaptation and stability of microvascular networks: theory and simulations. American Journal of Physiology-Heart and Circulatory Physiology 275 (2):H349–H360

33. Pries AR, Secomb TW (2008) Modeling structural adaptation of microcirculation. Microcirculation 15 (8):753–764

34. Pries AR, Secomb TW (2014) Making microvascular networks work: angiogenesis, remodeling, and pruning. Physiology 29 (6):446–455

35. Lindblom P, Gerhardt H, Liebner S, Abramsson A, Enge M, Hellström M, Bäckström G, Fredriksson S, Landegren U, Nyström HC (2003) Endothelial PDGF-B retention is required for proper investment of pericytes in the microvessel wall. Genes & development 17 (15):1835–1840

36. Newman AC, Nakatsu MN, Chou W, Gershon PD, Hughes CC (2011) The requirement for fibroblasts in angiogenesis: fibroblast-derived matrix proteins are essential for endothelial cell lumen formation. Molecular biology of the cell 22 (20):3791–3800

37. Zhang M, Zhao F, Zhang X, Brouwer LA, Burgess JK, Harmsen MC (2023) Fibroblasts alter the physical properties of dermal ECM-derived hydrogels to create a pro-angiogenic microenvironment. Materials Today Bio 23:100842

38. Rioja AY, Annamalai RT, Paris S, Putnam AJ, Stegemann JP (2016) Endothelial sprouting and network formation in collagen-and fibrin-based modular microbeads. Acta biomaterialia 29:33–41

39. Floryan M, Cambria E, Blazeski A, Coughlin MF, Wan Z, Offeddu G, Vinayak V, Kant A, Whisler J, Shenoy V (2025) Long-term physiological flow rescues regressed microvascular networks and increases their longevity. npj Biological Physics and Mechanics 2 (1):24

40. Wan Z, Zhang S, Zhong AX, Shelton SE, Campisi M, Sundararaman SK, Offeddu GS, Ko E, Ibrahim L, Coughlin MF (2021) A robust vasculogenic microfluidic model using human immortalized endothelial cells and Thy1 positive fibroblasts. Biomaterials 276:121032

41. Wan F, Chen F, Fan Y, Chen D (2022) Clinical Significance of TET2 in Female Cancers. Front Bioeng Biotechnol 10:790605. doi:10.3389/fbioe.2022.790605

42. Cao G, Xuan X, Hu J, Zhang R, Jin H, Dong H (2022) How vascular smooth muscle cell phenotype switching contributes to vascular disease. Cell Communication and Signaling 20 (1):180

43. Munn LL, Bazou D (2023) A Self-Assembly Method for Creating Vascularized Tumor Explants Using Biomaterials for 3D Culture. In: Cancer Cell Culture: Methods and Protocols. Springer, pp 211–220

44. Bazou D, Maimon N, Gruionu G, Munn LL (2016) Self-assembly of vascularized tissue to support tumor explants in vitro. Integr Biol (Camb). doi:10.1039/c6ib00108d

45. Shi Y, Meng J, Zhang S, Yang S, Liu G, Wu Z, Shen C, Shi C (2025) ERBB4 as a therapeutic target in aortic dissection: Implications for cell-based therapies in vascular regeneration. Biomol Biomed 25 (10):2324–2334. doi:10.17305/bb.2025.11925

46. Mistriotis P, Bajpai VK, Wang X, Rong N, Shahini A, Asmani M, Liang MS, Wang J, Lei P, Liu S, Zhao R, Andreadis ST (2017) NANOG Reverses the Myogenic Differentiation Potential of Senescent Stem Cells by Restoring ACTIN Filamentous Organization and SRF-Dependent Gene Expression. Stem Cells 35 (1):207–221. doi:10.1002/stem.2452

47. Chandra G, Defour A, Mamchoui K, Pandey K, Mishra S, Mouly V, Sreetama S, Mahad Ahmad M, Mahjneh I, Morizono H, Pattabiraman N, Menon AK, Jaiswal JK (2019) Dysregulated calcium homeostasis prevents plasma membrane repair in Anoctamin 5/TMEM16E-deficient patient muscle cells. Cell Death Discov 5:118. doi:10.1038/s41420-019-0197-z

48. Nguyen AT, Gomez D, Bell RD, Campbell JH, Clowes AW, Gabbiani G, Giachelli CM, Parmacek MS, Raines EW, Rusch NJ (2013) Smooth muscle cell plasticity: fact or fiction? Circulation research 112 (1):17–22

49. Sewell-Loftin MK, Bayer SVH, Crist E, Hughes T, Joison SM, Longmore GD, George SC (2017) Cancer-associated fibroblasts support vascular growth through mechanical force. Scientific reports 7 (1):12574

50. Erdogan B, Webb DJ (2017) Cancer-associated fibroblasts modulate growth factor signaling and extracellular matrix remodeling to regulate tumor metastasis. Biochemical Society Transactions 45 (1):229–236

51. Holter JC, Chang C-W, Avendano A, Garg AA, Verma AK, Charan M, Ahirwar DK, Ganju RK, Song JW (2022) Fibroblast-derived CXCL12 increases vascular permeability in a 3-D microfluidic model independent of extracellular matrix contractility. Frontiers in Bioengineering and Biotechnology 10:888431

52. Santi A, Kay EJ, Neilson LJ, McGarry L, Lilla S, Mullin M, Paul NR, Fercoq F, Koulouras G, Rodriguez Blanco G (2024) Cancer-associated fibroblasts produce matrix-bound vesicles that influence endothelial cell function. Science signaling 17 (827):eade0580

53. Bazou D, Maimon N, Gruionu G, Grahovac J, Seano G, Liu H, Evans CL, Munn LL (2018) Vascular beds maintain pancreatic tumour explants for ex vivo drug screening. Journal of tissue engineering and regenerative medicine 12 (1):e318–e322

54. Bondjers C, Kalén M, Hellström M, Scheidl SJ, Abramsson A, Renner O, Lindahl P, Cho H, Kehrl J, Betsholtz C (2003) Transcription profiling of platelet-derived growth factor-B-deficient mouse embryos identifies RGS5 as a novel marker for pericytes and vascular smooth muscle cells. The American journal of pathology 162 (3):721–729

55. Berger M, Bergers G, Arnold B, Hammerling GJ, Ganss R (2005) Regulator of G-protein signaling-5 induction in pericytes coincides with active vessel remodeling during neovascularization. Blood 105 (3):1094–1101

56. Naylor AJ, McGettrick HM, Maynard WD, May P, Barone F, Croft AP, Egginton S, Buckley CD (2014) A differential role for CD248 (Endosialin) in PDGF-mediated skeletal muscle angiogenesis. PloS one 9 (9):e107146

57. Hasanov Z, Ruckdeschel T, König C, Mogler C, Kapel SS, Korn C, Spegg C, Eichwald V, Wieland M, Appak S (2017) Endosialin promotes atherosclerosis through phenotypic remodeling of vascular smooth muscle cells. Arteriosclerosis, thrombosis, and vascular biology 37 (3):495–505

58. Liu S, Xu C, Zhang K, Han D, Yang F, Li Y, Zhao X, Ma S, Li H, Lu S (2021) CD248 as a bridge between angiogenesis and immunosuppression: a promising prognostic and therapeutic target for renal cell carcinoma. Annals of Translational Medicine 9 (23):1741

59. Naylor AJ, Azzam E, Smith S, Croft A, Poyser C, Duffield JS, Huso DL, Gay S, Ospelt C, Cooper MS (2012) The mesenchymal stem cell marker CD248 (endosialin) is a negative regulator of bone formation in mice. Arthritis & Rheumatism 64 (10):3334–3343

60. Ito S, Yokoyama U, Nakakoji T, Cooley MA, Sasaki T, Hatano S, Kato Y, Saito J, Nicho N, Iwasaki S (2020) Fibulin-1 integrates subendothelial extracellular matrices and contributes to anatomical closure of the ductus arteriosus. Arteriosclerosis, thrombosis, and vascular biology 40 (9):2212–2226

61. Liu G, Cooley M, Jarnicki A, Hsu A, Nair P, Haw T Fibulin-1 regulates the pathogenesis of tissue remodeling in respiratory diseases. JCI insight. 2016; 1 (9).

62. Ji T, Yan D, Huang Y, Luo M, Zhang Y, Xu T, Gao S, Zhang L, Ruan L, Zhang C (2024) Fibulin 1, targeted by microRNA-24-3p, promotes cell proliferation and migration in vascular smooth muscle cells, contributing to the development of atherosclerosis in APOE-/-mice. Gene 898:148129

63. Cooley MA, Kern CB, Fresco VM, Wessels A, Thompson RP, McQuinn TC, Twal WO, Mjaatvedt CH, Drake CJ, Argraves WS (2008) Fibulin-1 is required for morphogenesis of neural crest-derived structures. Developmental biology 319 (2):336–345

64. Argraves WS, Tran H, Burgess WH, Dickerson K (1990) Fibulin is an extracellular matrix and plasma glycoprotein with repeated domain structure. The Journal of cell biology 111 (6):3155–3164

65. Li Y, Zeng H, Zhong X (2025) Complement C3 in panvascular disease: a central integrator of immune signaling and vascular remodeling. Clinical Science 139 (21):1373–1403

66. Jensen LF, Bentzon JF, Albarrán-Juárez J (2021) The phenotypic responses of vascular smooth muscle cells exposed to mechanical cues. Cells 10 (9):2209

